# More than half of the variance in *in-vivo* ^1^H-MRS metabolite estimates is common to all metabolites

**DOI:** 10.1101/2022.09.29.510115

**Authors:** James J. Prisciandaro, Helge J. Zöllner, Saipavitra Murali-Manohar, Georg Oeltzschner, Richard A. E. Edden

## Abstract

The present study characterized associations among brain metabolite levels, applying bivariate and multivariate (i.e., factor analysis) statistical methods to tCr-referenced estimates of the major PRESS ^1^H-MRS metabolites (i.e., tNAA/tCr, tCho/tCr, mI/tCr, Glx/tCr), acquired from medial parietal lobe in a large (*n*=299), well-characterized international cohort of healthy volunteers (Povazan et al., 2020). Results supported the hypothesis that ^1^H-MRS-measured metabolite estimates are moderately intercorrelated (*M*_*r*_ = 0.42, *SD*_*r*_ = 0.11, *p*s < 0.001), with more than half (i.e., 57%) of the total variability in metabolite estimates common to (i.e., shared by) all metabolites. Older age was significantly associated with lower levels of common metabolite variance (CMV; *β* = -0.09, *p* = 0.048), despite *not* being associated with levels of any individual metabolite. Holding CMV levels constant, females had significantly lower levels of total choline (i.e., unique metabolite variance or UMV; *β* = -0.19, *p* < 0.001), mirroring significant bivariate correlations between sex and total choline reported previously. If replicated, these results would suggest that applied ^1^H-MRS researchers should shift their analytical framework from examining bivariate associations between individual metabolites and specialty-dependent (e.g., clinical, research) variables of interest (e.g., using t-tests) to examining multi-variable (i.e., covariate) associations between multiple metabolites and specialty-dependent variables of interest (e.g., using multiple regression). Without this shift, clear interpretation of associations of ^1^H-MRS metabolites with specialty-dependent variables of interest may not be possible.

## 1. Introduction

Proton MR spectroscopy (^1^H-MRS) is a widely used research and clinical diagnostic tool that combines standard MRI hardware with specialized pulse sequences to estimate the localized concentrations of products of cellular metabolism, or metabolites, *in vivo*. Most ^1^H-MRS data are acquired from brain tissue using single-voxel (i.e., one volume, or “voxel,” at a time) localization pulse sequences like STEAM, semi-LASER or PRESS, and implementations of these are included with most commercially available MRI systems (de Graaf, 2019). With the help of specialized post-processing (e.g., FID-A; Simpson et al., 2017) and linear-combination modeling (e.g., LCModel; Provencher, 1993) software, PRESS ^1^H-MRS in brain tissue at the commonest clinical MR field strengths (i.e., 1.5 T and 3 T) can reliably resolve five major metabolite signals from overlapping metabolite and nuisance signals (e.g., water, lipids): N-acetylaspartate plus N-acetylaspartylglutamate (i.e., total NAA or tNAA), glycerophosphocholine plus phosphocholine (i.e., total choline or tCho), creatine plus phosphocreatine (i.e., total creatine or tCr), myo-inositol (mI), and glutamate plus glutamine (Glx). The amplitude of each resolved metabolite signal is proportional to the number of signal-generating molecules in the voxel but is also influenced by idiographic characteristics of MRI hardware properties and the measurement process itself (e.g., coil loading, receiver gain, amplifier non-linearities). Signals are therefore typically normalized to an internal spectroscopic reference signal that is affected by the same characteristics (i.e., most often tCr or unsuppressed water) to allow inference across instruments (de Graaf, 2019). Once quantified, interpretation of metabolite variance typically relies on abstracting to accepted literature definitions of each metabolite: tNAA as a marker of neuronal viability or density, tCho of membrane/phospholipid turnover, tCr of bioenergetics (i.e., assuming tCr was not instead used as an internal reference), mI of osmotic regulation, and Glx of metabolic activity and/or neurotransmission (Rae, 2014).

There are many potential sources of inter-metabolite (i.e., metabolite-metabolite) covariance (e.g., shared biological pathways of synthesis and uptake, but also spectroscopic signal overlap, normalization, and differential likelihood of being present in particular tissue types). Inter-metabolite covariance has nonetheless been largely ignored in the ^1^H-MRS literature; we were unable, for example, to find a single published study characterizing associations among the major ^1^H-MRS metabolites in a reasonably large sample of healthy volunteers. In contrast, several application studies have reported bivariate correlations between theoretically selected pairs of metabolites and have generally demonstrated at least moderate inter-metabolite covariance (e.g., Glx and NAA: Kraguljac et al., 2012; Oeltzschner et al., 2019; Glx and GABA: Steel et al., 2020, He et al., 2021; GABA and mI: Oeltzschner et al., 2015). Furthermore, the influence of some specific sources of inter-metabolite covariance (e.g., normalization) has been well-characterized for similar applications of NMR spectroscopy, for example, in the nutritional and food sciences (Capozzi et al., 2011). Together, these lines of evidence provide support for the hypothesis that major ^1^H-MRS metabolites are moderately intercorrelated, due to some combination of substantive (e.g., common underlying biology) and methodological factors (e.g., shared localization in tissue, signal normalization, signal overlap).

Full characterization of inter-metabolite covariance should include analyses that go beyond estimating bivariate correlations between pairs of ^1^H-MRS metabolites, since bivariate correlations cannot identify and isolate sources of common (i.e., shared) metabolite variance (CMV) that affect more than two metabolite variables (e.g., normalization). These can only be determined and quantified using multivariate statistical methods, for example, factor analysis, an unsupervised learning algorithm that reduces the dimensionality of a metabolite dataset by identifying linear combinations of variables (or “factors”) that maximally explain observed inter-variable covariances (Watkins, 2018). Such multivariate methods have been productively used in NMR research for over forty years (Malinowski, 1978), and have been recommended for ^1^H-MRS research and clinical applications (El-Deredy, 1997), but have not been widely adopted to date. Only a few studies that we are aware of, each conducted in relatively small clinical samples (i.e., individuals with HIV-associated dementia; Yiannoutsos et al., 2004; Mohamed et al., 2010), have used factor analysis to identify and isolate CMV factors (e.g., “inflammation,” marked by CMV in Cho/Cr and mI/Cr across multiple brain regions).

Since multiple-regression research has shown that observed bivariate associations are often spurious results of shared covariance with confounding ‘third variable(s)’ (Cohen et al., 2002), it is imperative to not only obtain robust bivariate and multivariate estimates of inter-metabolite covariance in ^1^H-MRS data, but to also evaluate the extent to which associations of specialty-dependent (e.g., clinical, behavioral) variables of interest with individual metabolites are driven by CMV versus variance unique to those individual metabolites (i.e., unique metabolite variance, UMV), for example, using statistical covariate analyses (e.g., ANCOVA, multiple regression). Unfortunately, the vast majority of applied ^1^H-MRS studies to date have only evaluated bivariate associations of one or a few theoretically selected metabolites with specialty-dependent variables of interest, ignoring the potential influence of inter-metabolite covariance on these associations. As a result, it is possible that some, many, most, or all reported associations between ^1^H-MRS metabolites and specialty-dependent variables of interest have been driven by CMV, thereby undermining authors’ interpretations of these associations as reflecting something particular about their chosen ^1^H-MRS metabolites (i.e., UMV).

The primary goal of this study was to characterize inter-metabolite covariance in a large (*n*=299), well-characterized international cohort of healthy volunteers (Povazan et al., 2020), applying both bivariate and multivariate (i.e., factor analysis) statistical methods to tCr-normalized estimates of the major PRESS ^1^H-MRS metabolites (tNAA/tCr, tCho/tCr, mI/tCr, Glx/tCr). The secondary goal of this study was to evaluate whether observed associations of the major metabolites with available covariates (i.e., age, sex) were driven by CMV and/or UMV in the same cohort of volunteers.

## 2. Material and methods

### 2.1 Participants and data (acquisition, preprocessing, and modeling)

Data for the present study, along with acquisition details, have been published previously (Povazan et. al., 2020); as noted therein, institutional review board approval and participant written informed consent were obtained at each study site. Briefly, 299 healthy volunteers from 26 sites (mean [SD] *n* per site = 11.19 [1.80]) contributed ^1^H-MRS data from 3×3×3 cm^3^ medial parietal lobe voxels (*Figure 1a*), acquired using vendor-native short-TE PRESS sequence on scanners from 3 vendors (Philips: 10 sites, total *n* = 119; Siemens: 8 sites, total *n* = 89; GE: 8 sites, total *n* = 91), with the following parameters: TR/TE = 2000/35 ms; 64 averages; 2, 4, or 5 kHz spectral bandwidth; 2048-4096 data points; 140, 50, or 150 Hz water suppression pulse bandwidth for Philips, Siemens, or GE, respectively; acquisition time = 2.13 min. Eight to sixteen averages of water reference signal were acquired with similar parameters, but without water suppression. Institutional review board approval and participant written informed consent were obtained at each study site. Data were preprocessed following expert-consensus-recommended practices (e.g., frequency and phase correction of individual transients in the time domain [Mikkelsen et al., 2020; Near et al., 2021], eddy current correction [Barkhuijsen et al., 1987, and HSVD water removal [Klose, 1990]) using the open-source MATLAB toolbox Osprey (Oeltzschner et al., 2020). Vendor-specific basis sets were created for linear-combination modeling using the MATLAB toolbox FID-A (Simpson et al., 2017) as detailed in Zollner et al., 2021. The LCModel version 6.3 algorithm (Provencher, 1993) was used to fit the pre-processed data with these basis sets and default settings (*Figure 1b*). Quantitative metabolite estimates were derived from the LCModel-generated model amplitudes in Osprey and subsequently referenced to (i.e., divided by) estimated tCr levels (Povazan et al., 2020), resulting in four metabolite variables (i.e., tNAA/tCr, tCho/tCr, mI/tCr, and Glx/tCr) for further statistical analyses. Data quality metrics (signal-to-noise (SNR) and linewidth of the NAA singlet) were calculated for the pre-processed data. SNR was defined as the ratio of the peak height and the standard deviation of the frequency-domain spectrum between -2 and 0 ppm. Linewidth was defined as the average of the simple FWHM and the FWHM of a Lorentzian peak model. A summary table following the minimum reporting standards recommended by a recent MRS consensus paper (Lin et al., 2021) can be found in *Supplementary Table 1*.

**Figure 1.**
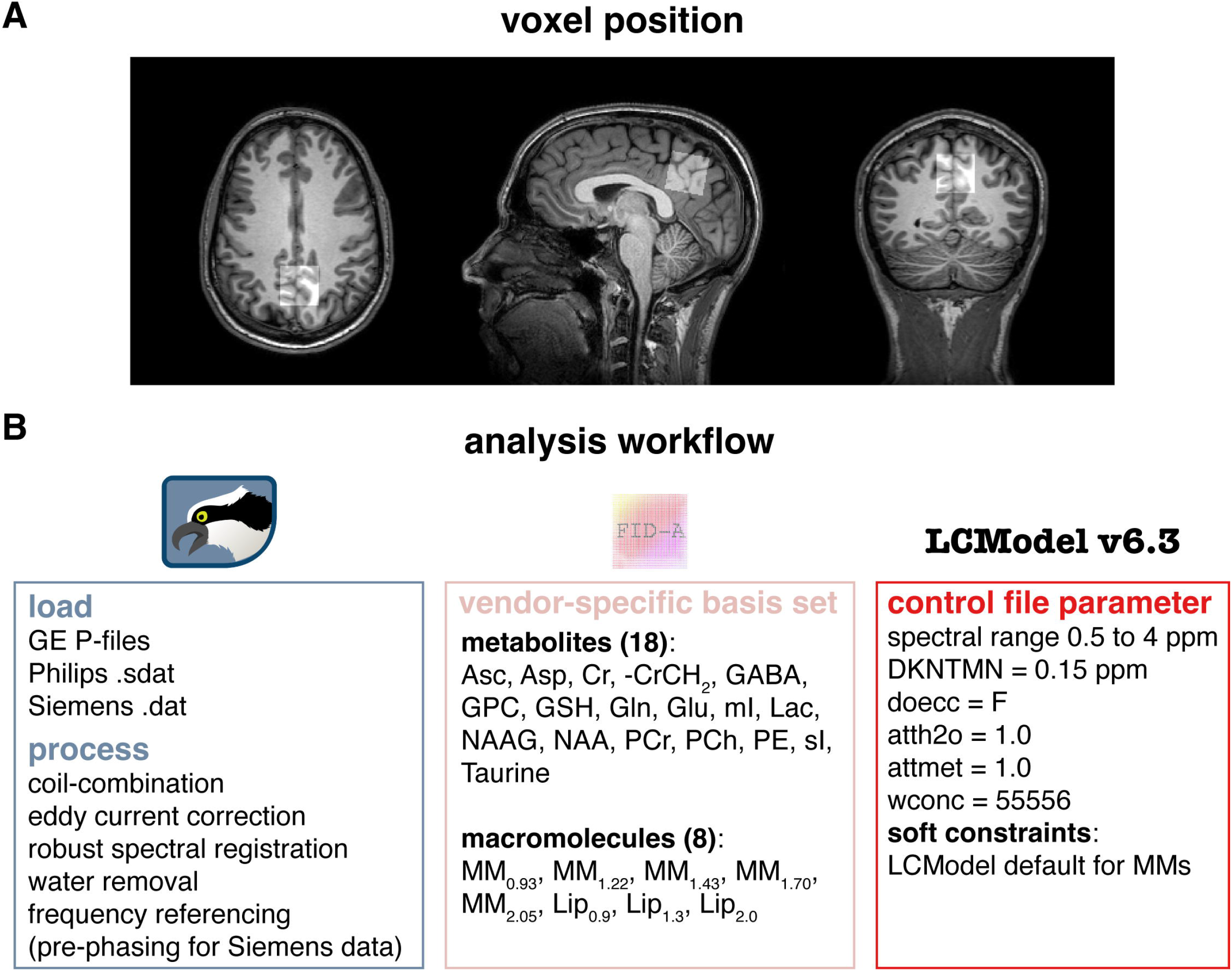
*a*. Sample 3×3×3 cm^3^ medial parietal lobe voxel position. *b*. Detailed schematic of the analysis workflow used in this study.

### 2.2 Statistical analyses

Statistical analyses were performed on group- (i.e., site-) mean centered metabolite variables (i.e., to provide clearly interpretable and unbiased estimates of inter-metabolite covariance; Enders & Tofighi, 2007) in Mplus version 8.7 software (Muthen et al., 2017), using a maximum likelihood estimator with robust standard errors (MLR) and with “type=complex” specified to adjust model-fit indices and standard errors for the complex sampling design of the data (i.e., with participants clustered by site [“cluster = site”] and sites stratified by vendor [“stratification = vendor”]; Muthen et al., 1995; Stapleton, 2008).

Pearson correlation coefficients were calculated for each pair of metabolite variables, as well as for each available combination of covariate and metabolite variable (e.g., age and Glx/tCr); for correlation coefficients involving the binary variable, sex, resulting values were point-biserial coefficients with p-values equal to those derived from equivalent independent samples t-tests (i.e., with sex as the independent, and metabolite as the dependent, variable) (Cohen et al., 2002).

Exploratory factor analysis (EFA) was conducted to obtain eigenvalues associated with each extracted factor, representing the total amount of CMV explained by a given factor, with eigenvalues < 1 typically interpreted as denoting ignorable factors (Watkins, 2018). Only a single CMV factor was ultimately modeled using confirmatory factor analysis (CFA), because the observed data consisted of only 4 dependent variables, and each modeled factor in a factor analysis should feature ≥ 3 highly associated (or “loading”) dependent variables to avoid estimation problems and to ensure model reliability (Watkins, 2018). Factor variances were fixed to “1” (i.e., standardized) for model identification, allowing all factor loadings (i.e., metabolite on factor regression coefficients) to be freely estimated (Watkins, 2018). Correlations among model residuals were fixed to “0” to impart the standard, conservative baseline assumption of local independence (Henning, 1989). Goodness of model fit to the data was evaluated using the comparative fit index (CFI, cutoff ≥ 0.95), Tucker Lewis index (TLI, cutoff ≥ 0.95), and standardized root mean square residual (SRMR, cutoff ≤ 0.08) (Hooper et al., 2008; Taasoobshirazi et al., 2016). Modification indices (MIs) were generated for constrained model parameters (i.e., correlations among model residuals), reflecting the improvement in model fit (x^2^) that would be achieved if a given constrained parameter were to be freely estimated. Constraints associated with very high MIs (i.e., > 10, corresponding to *p*-values ≤ 0.001) were considered for free estimation, one at a time.

From the final CFA model, a Multiple Indicator Multiple Causes (MIMIC; Joreskog & Sorbom, 1996) model was constructed to evaluate associations of age and sex with the identified CMV factor and each of its constituent metabolite variables. The standard, stepwise forward modeling procedure was used to construct the final MIMIC model (Brown, 2006); first regressing the CMV factor on covariates (i.e., while simultaneously constraining each regression path from covariates to individual metabolite variables to “0”), then using MIs to unconstrain only those paths that would result in significant improvement to model fit if freely estimated (i.e., > 10, corresponding to p-values ≤ 0.001), one at a time. Regression paths unconstrained using this method, if any, represent associations of covariates with individual metabolites that are statistically significant at equal levels of (i.e., statistically controlling for) the CMV factor, allowing for dissociation of covariate effects driven by CMV (i.e., CMV factor regressed on covariates) versus UMV (i.e., individual metabolite variables simultaneously regressed on covariates).

Fully standardized model parameters are provided for final CFA and MIMIC models, including factor loadings, estimated covariate regression coefficients, UMV (i.e., variance in each metabolite unaccounted for by the CMV factor and/or covariates, along with its reciprocal [i.e., *r*^*2*^]), and estimated residual correlation coefficients, if any.

## 3. Results

### 3.1 Participants and data acquisition, preprocessing, and modeling

Data were generally of excellent quality (NAA SNR: *M* = 213.12, *SD* = 68.74; Water FWHM: *M* = 6.83 Hz, *SD* = 0.79 Hz). However, data from 8 participants were excluded after visual inspection of the modeling outputs due to incorrectly placed voxels (*n* = 2, vendor = Siemens), unacceptably high lipid peaks (*n* = 4, vendor = Siemens; *n* = 1, vendor = Philips), or frequency and phase correction failure (*n* = 1, vendor = GE). Data from an additional 3 participants were excluded following metabolite quantification because their tNAA/tCr (n = 1, vendor = GE) or Glx/tCr (n = 1, vendor = GE; n = 1, vendor = Philips) values were identified as extreme outliers (i.e., ≥ 3^rd^ quartile + 3 * interquartile range), leaving 288 participants’ data available for covariation analyses. *Figure 2* shows mean data, model, and residual, while *Supplementary Figure 1* shows participant data, model, and residual. LCModel fits of these data were excellent, as indicated by their Cramer Rao Lower Bound (CRLB), a measure of modeling uncertainty (tNAA *M* = 1.41 %, *SD* = 0.50 %; tCho *M* = 2.67 %, *SD* = 1.00 %; tCr *M* = 1.36 %, *SD* = 0.53 %, mI *M* = 3.76 %, *SD* = 0.65 %; Glx *M* = 4.61 %, *SD* = 0.67 %).

**Figure 2.**
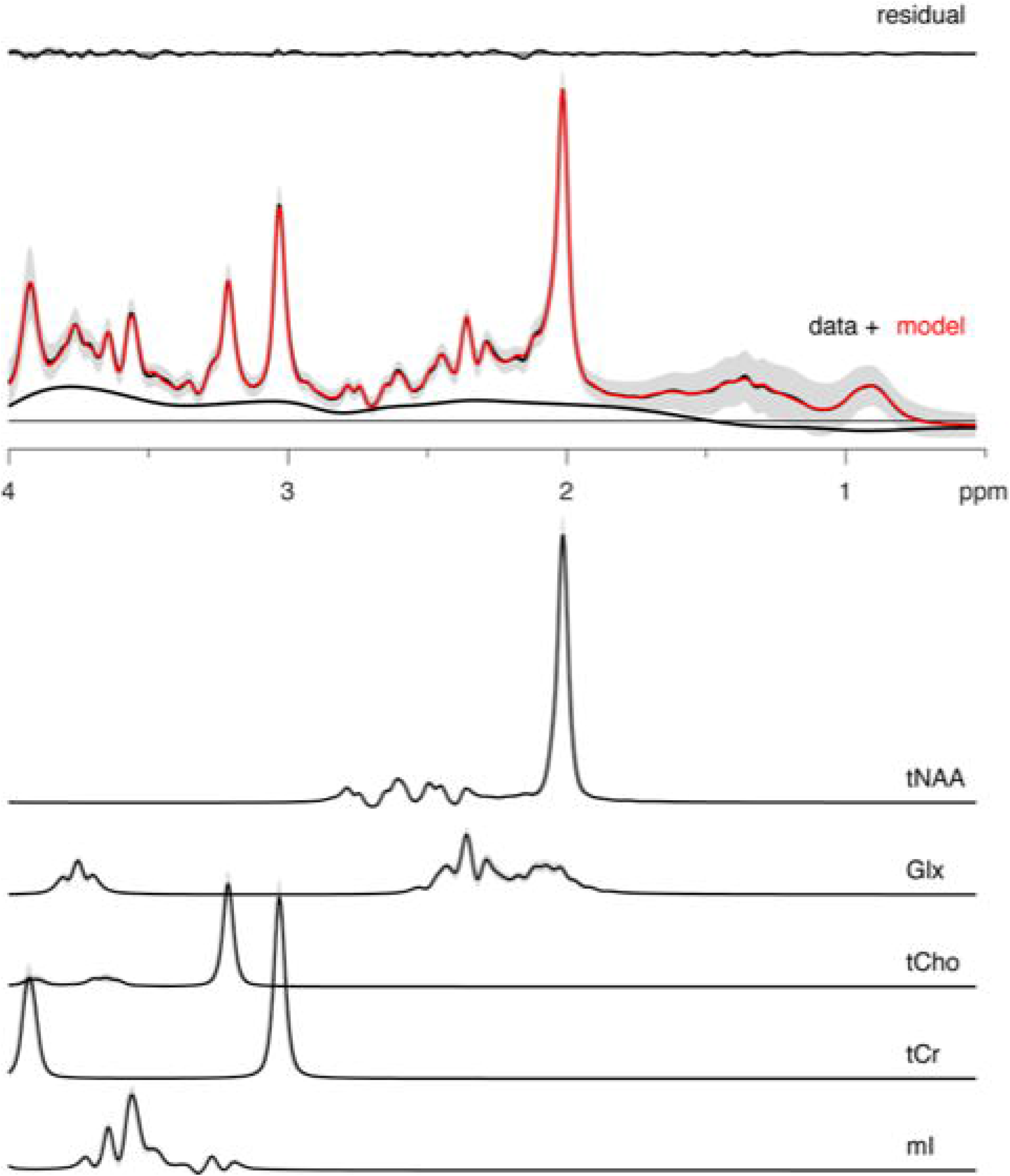
Mean spectra (black) +/ SD (gray ribbons) and mean fit (red); mean residual (above) and mean contribution from individual metabolites (i.e., tNAA total NAA, Glx glutamate+glutamine, tCho total choline, tCr total creatine, mI myo-inositol) +/- SD (gray ribbons) (below).

### 3.2 Statistical analyses

*Figure 3* shows scatterplots and linear regression lines, along with Pearson correlation coefficients, for each pair of tCr-referenced metabolite variables. Inter-metabolite correlations were generally moderate in strength (*M*_*r*_ = 0.42, *SD*_*r*_ = 0.11) and were uniformly positive and significantly different from zero (*p*s < 0.001).

**Figure 3.**
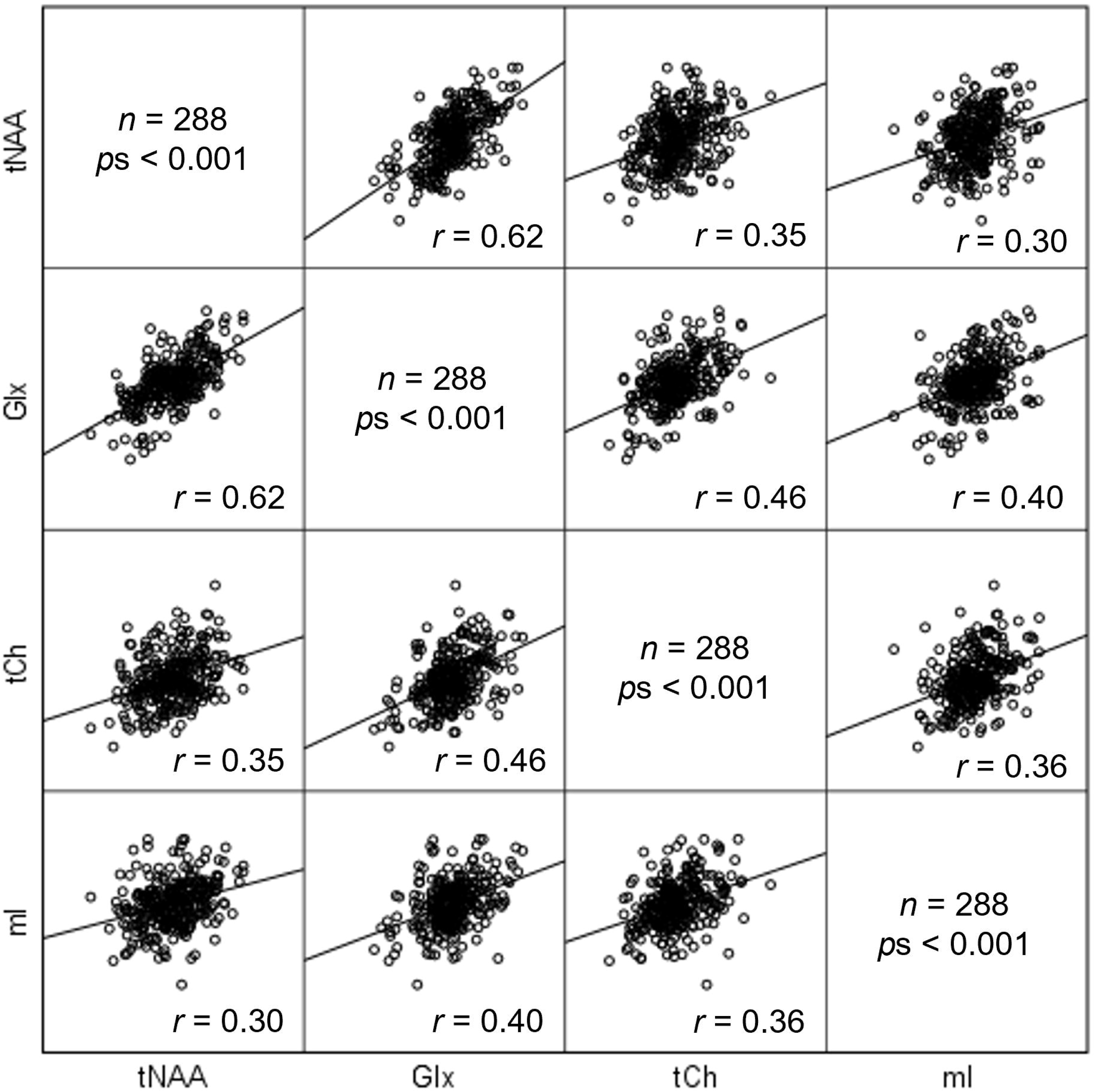
Scatterplot matrix, featuring linear fit lines, depicting bivariate associations between each pair of metabolites (i.e., tNAA total NAA, Glx glutamate+glutamine, tCho total choline, tCr total creatine, mI myo-inositol).

EFA-estimated eigenvalues (1: 2.26, 2: 0.75, 3: 0.63, 4: 0.36) strongly supported extraction of a single factor that explained 57% of total metabolite variance, and models with > 1 extracted factor failed to converge, as predicted. As a result, a single factor CFA model was constructed that provided good (CFI = 0.98, SRMR = 0.03) to adequate (TLI = 0.93) fit to the data; no MIs exceeded the pre-defined threshold of *p* ≤ 0.001, and so the baseline assumption of local independence (i.e., that the model fully explained all inter-metabolite covariances) was retained. *Figure 4* shows a diagram of the final CFA model, labeled by its standardized parameter estimates and standard errors. Factor loadings were moderate-strong (M_*β*_ = 0.64, SD_*β*_ = 0.18) and universally statistically significant (*p*s < 0.001), with the CMV factor explaining between 22% (mI/tCr) and 76% (Glx/tCr) of the variance in each metabolite (*M*_*r*_^*2*^ = 0.56, *SD*_*r*_^*2*^ = 0.24).

**Figure 4.**
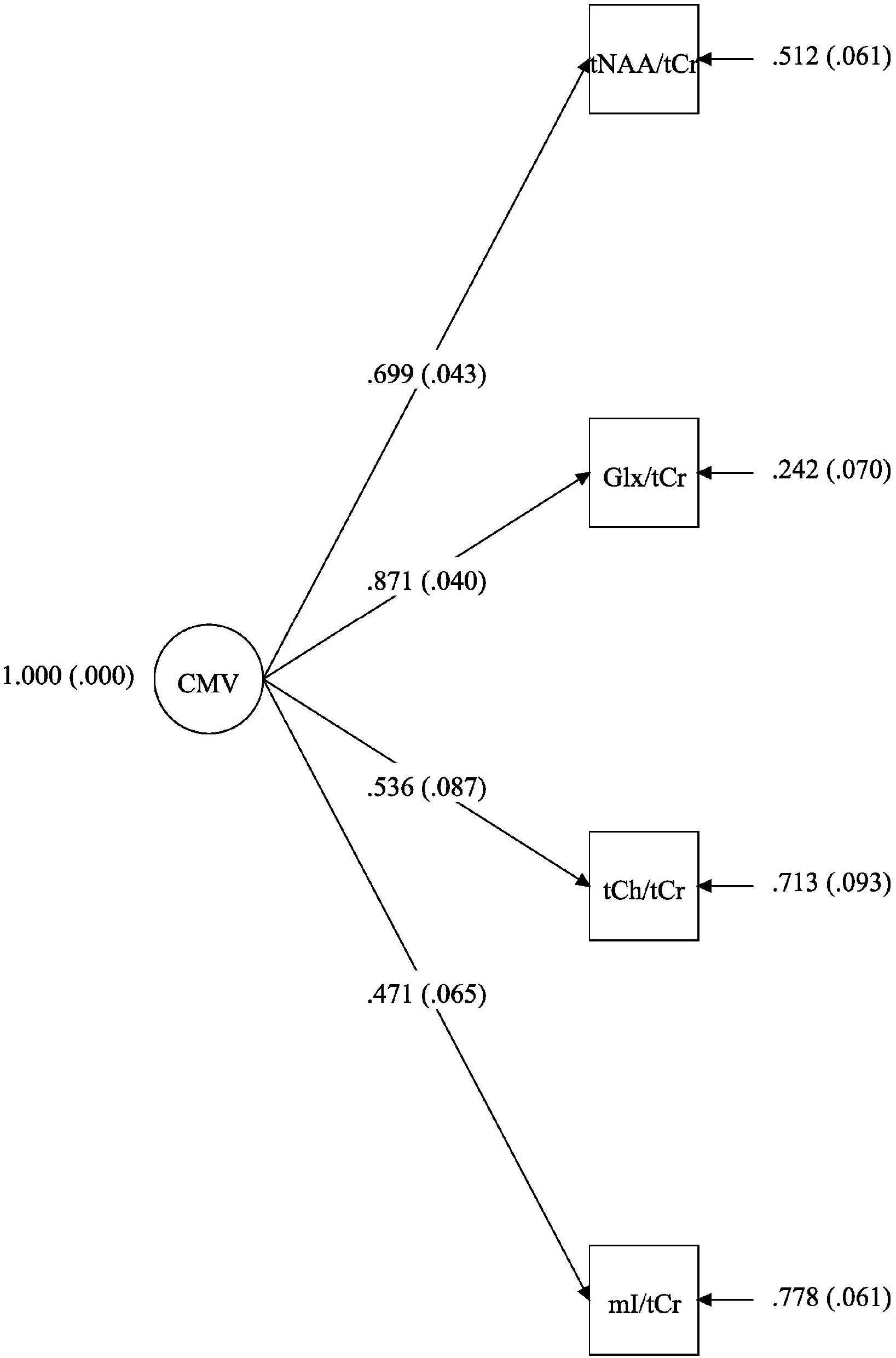
Final confirmatory factor analysis model, labeled with fully standardized (ranging between “0” and “1”) parameter estimates (standard errors). Parameter estimates on arrowed lines from the common metabolite variance (CMV) factor to metabolite variables represent the strength of the relationship between each metabolite and the CMV factor. Parameter estimates adjacent to arrowed lines pointing to each metabolite represent the unique metabolite variance (UMV) that was not explained by the CMV factor in that metabolite (and, its inverse, the amount of variance that was explained by the CMV factor). tNAA = total NAA, Glx = glutamate+glutamine, tCho = total choline, tCr = total creatine, mI = myo-inositol.

Age (*M* = 26.46, *SD* = 4.62) and sex (female *n* [%] = 147 [51%]) were significantly associated with one another (*r* = 0.12, *p* = 0.049). With the exception of sex with tCho/tCr (*r* = - 0.23, *p* < 0.001), bivariate associations of covariates with metabolite variables were small (age *M*[*SD*]_*r*_ = -0.04[0.05], sex *M*[*SD*]_*r*_ = -0.05[0.05]) and non-significant (age *p*s > 0.192, sex *p*s > 0.11). The baseline MIMIC model, with the CMV factor regressed on age and sex but with all paths from covariates to metabolite variables constrained to “0,” provided mediocre (CFI = 0.92, SRMR = 0.05) to inadequate (TLI = 0.86) fit to the data. The constrained path from sex to tCho/tCr was associated with the largest, suprathreshold MI value (i.e., 13.95), supporting removal of the constraint and re-estimation of the model. Following this modification, one additional suprathreshold MI value (i.e., 10.38) was provided, supporting removal of the constrained residual correlation between tCho/tCr and mI/tCr (i.e., relaxing the assumption of local independence for these two variables). The final MIMIC model, which included these two modifications to the baseline model, provided excellent fit to the data (CFI/TLI = 1.00, SRMR = 0.02). *Figure 5* shows a diagram of the final MIMIC model, labeled by its standardized parameter estimates and standard errors. The regression path from age (β = -0.09, *p* = 0.048), but not sex (β = -0.11, *p* = 0.066), to the CMV factor was statistically significant. However, the variance explained in the CMV factor by age was small (i.e., 2%), likely owing to the restricted age range of the sample (i.e., 18-35 years). The freed regression path from sex to tCho/tCr was statistically significant (β = -0.19, *p* < 0.001), as anticipated by its associated suprathreshold MI value.

**Figure 5.**
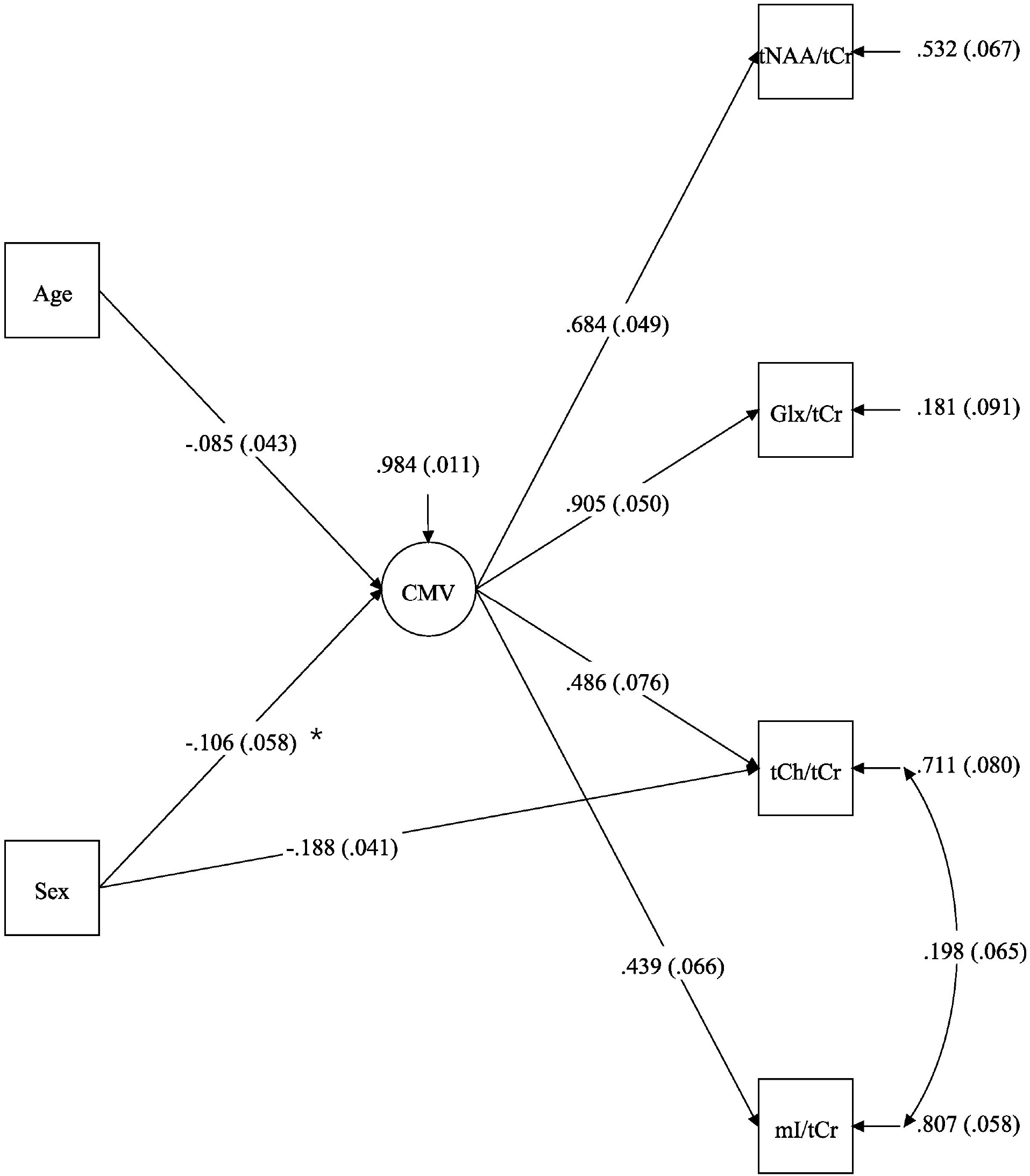
Final Multiple Indicator Multiple Causes (MIMIC) model, labeled with fully standardized parameter estimates (standard errors). Parameter estimates on top of single arrowed lines are regression coefficients representing the strength of the relationship between a given pair of variables or factors, while those on top of double arrowed lines are correlation coefficients representing the strength of the relationship between the unexplained variances of a given pair of metabolites. Parameter estimates adjacent to arrowed lines pointing to each metabolite or factor represent the variance in that metabolite or factor that was not explained by the model. Sex was coded, 0 = Male, 1 = Female. CMV = common metabolite variance, tNAA = total NAA, Glx = glutamate+glutamine, tCho = total choline, tCr = total creatine, mI = myo-inositol.

In summary, bivariate correlations among metabolite variables were generally moderate in strength and uniformly positive and significantly different from zero. A single factor CFA model provided good fit to the data, with the CMV factor explaining 57% of the variance in each metabolite, on average. MIMIC modeling found that older participants had lower levels of CMV, and that, controlling for CMV levels, females had lower levels of total choline (i.e., tCho/tCr) than males.

## 4. Discussion

The present study characterized associations among brain metabolite levels, applying bivariate and multivariate (i.e., factor analysis) statistical methods to tCr-referenced estimates of the major PRESS ^1^H-MRS metabolites (i.e., tNAA/tCr, tCho/tCr, mI/tCr, Glx/tCr), acquired from medial parietal lobe in a large (*n*=288), well-characterized international cohort of healthy volunteers (Povazan et al., 2020). Results supported the hypothesis that ^1^H-MRS-measured metabolite estimates are moderately intercorrelated, with more than half of the total variability in metabolite estimates common to (i.e., shared by) all metabolites. Older age was significantly associated with lower levels of common metabolite variance (CMV), despite *not* being associated with levels of any individual metabolite. Holding CMV levels constant, females had significantly lower levels of total choline (i.e., unique metabolite variance or UMV), mirroring the significant bivariate correlation between sex and total choline reported previously (Povazan et al., 2020). If replicated, these findings have broad and substantial implications for the analysis and interpretation of ^1^H-MRS data in applied studies.

Though both the present study and several applied studies (e.g., Kraguljac et al., 2012; Oeltzschner et al., 2019) have demonstrated that ^1^H-MRS acquired metabolite estimates are moderately intercorrelated, the vast majority of ^1^H-MRS applications have focused on one or a few theoretically selected metabolite estimates and their bivariate associations with specialty-dependent (e.g., clinical, behavioral) variables of interest, ignoring the potential influence of inter-metabolite covariance on these associations in their analyses and interpretations. If, however, as the results from the present study suggest, more than half of the total metabolite variance is common to all metabolites, such interpretations can only be valid if the associations are driven solely by the other half of the variance in the selected metabolite(s). Because most applied studies have only evaluated one or a few metabolites using bivariate statistical tests, it is possible that some, many, or most reported associations between ^1^H-MRS metabolites and specialty-dependent variables of interest have been driven by CMV. Results from the present study suggest that applied researchers broaden their analytical approach to include more metabolites, statistically controlling for CMV using appropriate covariance modeling methods (e.g., multiple regression) in place of more commonly used bivariate methods (e.g., t-test).

What exactly *is* this CMV factor that appears to account for over half of the variance in PRESS ^1^H-MRS metabolite estimates? Does it represent one or more common underlying biological and/or methodological phenomena? The first common factor extracted from biological data is typically interpreted as a measure of the overall biological process (e.g., “cellular metabolism,” in the case of ^1^H-MRS data), with the relative strength of associations between this factor and its constituent metabolites providing additional interpretational clues (Ripley, 1996). For example, although the CMV factor was significantly associated with each metabolite in the present study, it explained 3-4x more variance in Glx/tCr versus mI/tCr (with tNAA/tCr and tCho/tCr intermediary), and unlike with Glx/tCr and tNAA/tCr, the CMV factor did not adequately explain the covariance between mI/tCr and tCho/tCr. Based on this pattern of associations, one could hypothesize that the observed CMV factor reflects more neuronal metabolism than glial metabolism (Rae, 2014).

Because common factors are empirically derived, however, their meaning can arguably only be ascertained through the process of “construct validation” (Cronbach et al., 1955), whereby opposing predictions concerning the nature of CMV are systematically tested. For example, one way to determine whether CMV reflects biological versus methodological phenomena is to evaluate associations of CMV with substantive biological variables of interest (e.g., disease, age). In the present study, older age was significantly associated with lower levels of CMV, lending preliminary support to the hypothesis that CMV reflects biological processes, at least in part. Unfortunately, however, because participants in the present study fell between a restricted age range of 18-35 years (Povazan, 2020), age was only able to predict a small amount of CMV (approximately 2%). Future studies featuring less restricted samples will be necessary to determine the real relationship between age and CMV. In contrast, sex, which was not restricted in the present sample, was *not* associated with CMV, but was significantly associated with tCho/tCr UMV. Overall, though not definitive by any means, these preliminary construct validity findings demonstrate the power of the statistical approach taken in the present study to distinguish between associations of specialty-dependent variables of interest with CMV versus UMV.

Another way to determine whether CMV reflects methodological versus biological phenomena is to examine its generalizability across MR field strengths and ^1^H-MRS acquisition methods. For example, if CMV reflects shared metabolite variance due to the relatively poor resolvability of metabolite signals at common clinical field strengths (1.5 or 3 T), one would predict it to be substantially reduced in metabolite data acquired at higher field strengths (e.g., 7 T) or with advanced acquisition techniques designed to minimize overlap (e.g., J-difference editing, Rothman et al., 1993; two-dimensional J-resolved PRESS, Ryner et al., 1995). These techniques have the added advantage of potentially providing more metabolite variables for factor analyses, allowing for evaluation of more complicated models with multiple factors (Watkins, 2018). Luckily, much of the work needed to replicate and validate the CMV factor found in the present study can be done using data that already exists. Authors of published applied ^1^H-MRS studies could “kill two birds with one stone” by separating CMV from UMV in their published data and then examining whether their reported bivariate associations were indeed driven by UMV (i.e., in accordance with their original published interpretation) or CMV, thereby simultaneously evaluating the validity of the CMV factor and disambiguating interpretation of their published findings. Analyses for such re-examinations could be accomplished either directly, within the factor model itself (i.e., as was done in the present study) using specialized software like MPlus (Muthen et al., 2017) or the lavaan R toolbox (Rosseel, 2012), or indirectly, by first calculating factor and residual scores (i.e., the latter reflecting metabolite variance not attributable to the common factor plus random error) for each participant using software like IBM SPSS Statistics [IBM Corp., Armonk, N.Y., USA] or the psych R toolbox [Revelle, 2022].

In conclusion, the present study demonstrated in a large sample of healthy volunteers that PRESS ^1^H-MRS metabolite measures are moderately intercorrelated and that over half of the variance contained in these measures is CMV, common to all metabolites. Preliminary construct validity results further demonstrated that older age was associated with lower CMV levels, and that controlling for CMV levels, female sex was associated with lower mI/tCr UMV levels. If replicated, these results would suggest that applied ^1^H-MRS researchers should shift their analytical framework from examining bivariate associations between individual metabolites and specialty-dependent (e.g., clinical) variables of interest (e.g., using t-tests) to examining multi-variable (i.e., covariate) associations between multiple metabolites and specialty-dependent variables of interest (e.g., using multiple regression). Without this shift, clear interpretation of associations of ^1^H-MRS metabolites with specialty-dependent variables of interest may not be possible.

## Data availability statement

The de-identified neuroimaging data and related materials are available publicly through NITRC at the following URL: https://www.nitrc.org/projects/biggaba/

## List of abbreviations

CFA: confirmatory factor analysis
CFI: comparative fit index
CMV: common metabolite variance
EFA: exploratory factor analysis
Glx: glutamate plus glutamine
MI: modification indices
mI: myo-Inositol
MIMIC: Multiple Indicators Multiple Causes
MLR: maximum likelihood estimator with robust standard errors
SRMR: standardized root mean square residual
tCho: glycerophosphocholine plus phosphocholine
tCr: creatine plus phosphocreatine
TLI: Tucker Lewis Index
tNAA: N-acetylaspartate plus N-acetylaspartylglutamate
UMV: unique metabolite variance

