## Supplementary for "More than half of the variance in *in-vivo* ^1^H-MRS metabolite estimates is common to all metabolites"

**Supplementary Materials**

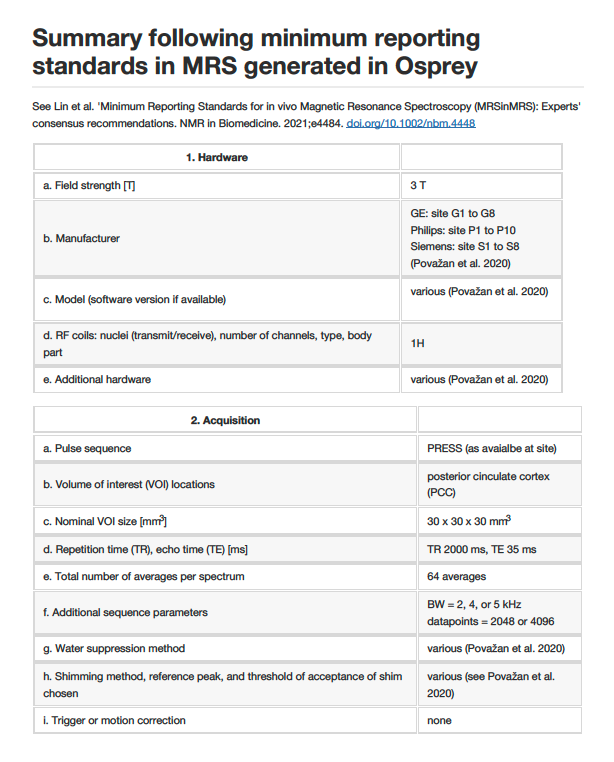

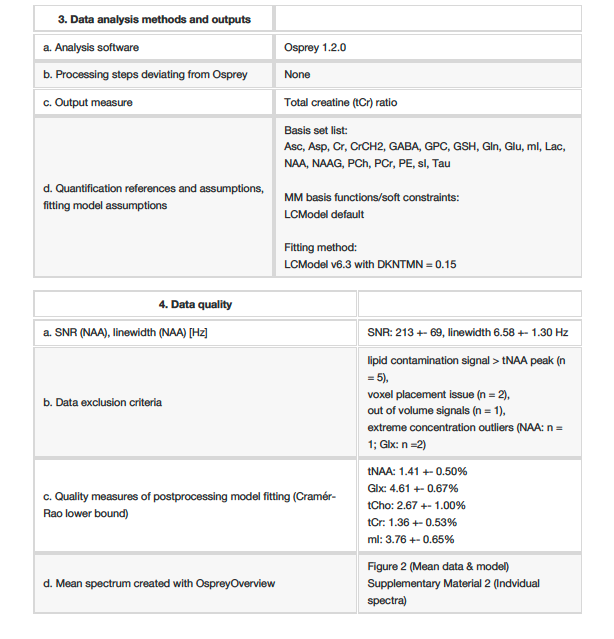

Supplementary Table 1 – MRSinMRS summary generated in Osprey

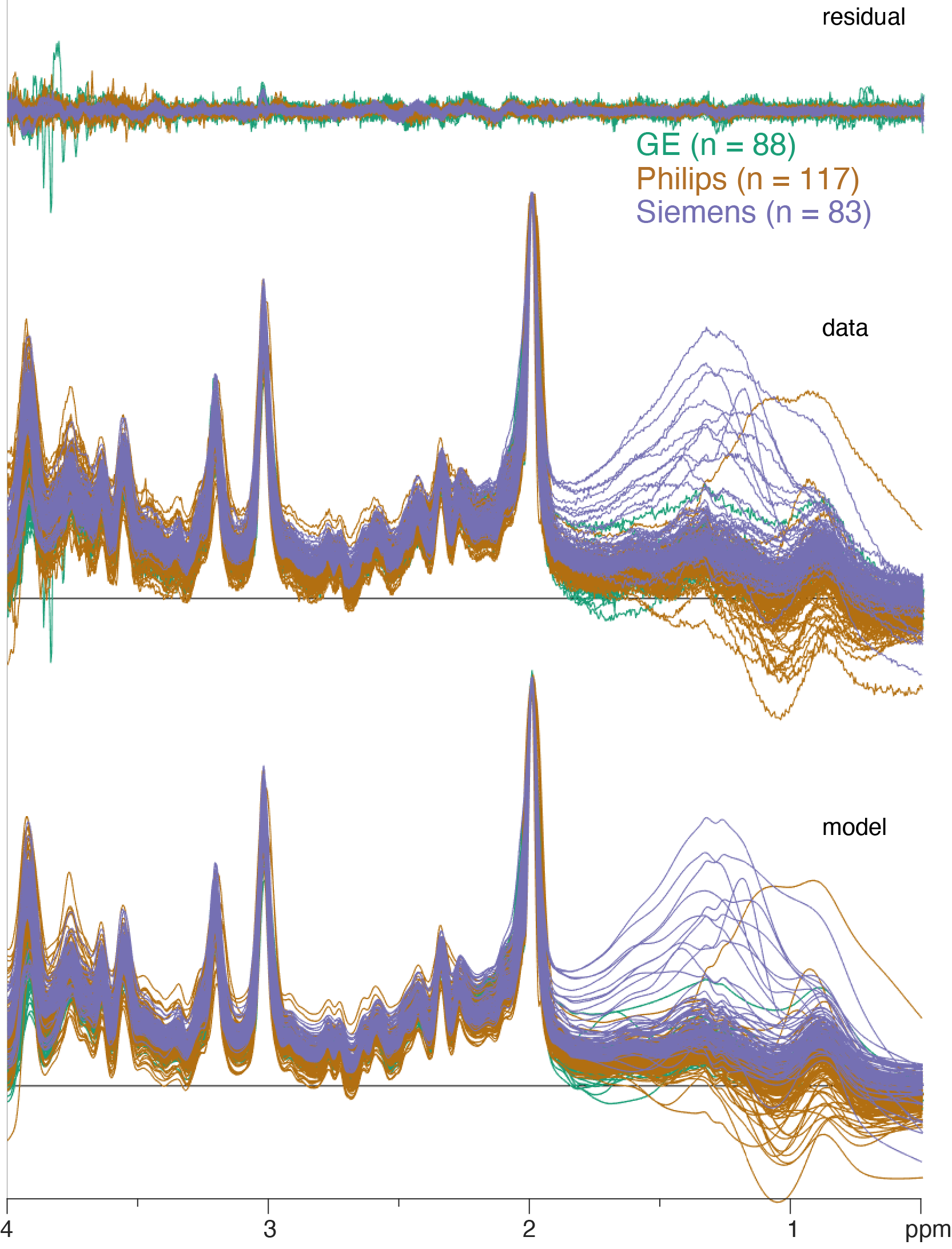

Supplementary Figure 2 – Overview of modeling results including all subjects.
